# Sexual selection matters in genetic rescue, but productivity benefits fade over time; a multi-generation experiment to inform conservation

**DOI:** 10.1101/2024.10.03.616221

**Authors:** George West, Michael Pointer, Will Nash, Rebecca Lewis, Matt J.G. Gage, David S. Richardson

**Affiliations:** School of Biological Sciences, University of East Anglia, Norwich Research Park, Norfolk, NR4 7TJ, UK; Earlham Institute, Norwich Research Park, Norfolk, NR4 7UZ, UK

**Keywords:** Genetic rescue, Sexual selection, Inbreeding depression, Small populations, Genetic variation, *Tribolium*

## Abstract

Globally, many species are threatened by population decline because of anthropogenic changes which leads to population fragmentation, genetic isolation, and inbreeding depression. Genetic rescue, the controlled introduction of genetic variation, is a method used to potentially relieve such effects in small populations. However, without understanding how the characteristics of rescuers impact rescue attempts interventions run the risk of being sub-optimal, or even counterproductive. We use the Red Flour Beetle (*Tribolium castaneum*) to test the impact of rescuer sex, and sexual selection background, on population productivity. We record the impact of genetic rescue on population productivity in 24/36 replicated populations for ten generations following intervention. We find little or no impact of rescuer sex on the efficacy of rescue but show that a background of elevated sexual selection makes individuals more effective rescuers. In both experiments, rescue effects diminish 6-10 generations after the rescue. Our results confirm that the efficacy of genetic rescue can be influenced by characteristics of the rescuers and that the level of sexual selection in the rescuing population is an important factor. We show that any increase in fitness associated with rescue may last for a limited number of generations, suggesting implications for conservation policy and practice.

## Introduction

Populations worldwide increasingly face extinction after becoming fragmented by human activity (Ceballos and Ehrlich, 2023). Fragmentation reduces population size and increases risk of genetic isolation, leading to increased impact of genetic drift and loss of genetic variation. Consequentially, many small populations suffer inbreeding depression (reduction in fitness when recessive, deleterious alleles appear in homozygous form, and/or the loss of heterozygote advantage) and reduced adaptive potential (Charlesworth and Willis, 2009; Crnokrak and Roff, 1998). Individuals within such populations are also more prone to environmental stress, which can exacerbate inbreeding depression (Frankham, 2005; Reed et al., 2002; Richardson et al., 2004). Interaction between these factors can lead to population or species extinction (Soule and Gilpin, 1986; Blomqvist et al., 2010; Palomares et al., 2012).

Genetic rescue, increasing population fitness through the introduction of novel alleles beyond the demographic effects of immigration, is one way to relieve inbreeding depression (Ingvarsson, 2001; Hedrick et al., 2011). This requires the introduction of rescuers (conspecific individuals from a different population), allowing reproduction with the inbred population. The aim is to introduce new genetic diversity, reducing homozygosity and the expression of deleterious alleles in offspring. Introducing genetic variation also increases adaptive potential, providing standing variation for selection to act on (Hoelzel et al., 2019; Mable, 2019) and increasing the potential for evolutionary rescue (Bell and Gonzalez, 2009; Schiffers et al., 2013; Lindsey et al., 2013).

Genetic rescue has been studied in wild, captive, and laboratory populations across many taxa (reviewed in Frankham, 2015, 2016; White et al., 2023) and has seen many successful implementations (Miller et al., 2020; Davis et al., 2021; Robinson et al., 2017; Pregler et al., 2022; Pavlova et al., 2023). Reviews and meta-analyses support its utility as a conservation tool (Waller, 2015; Whiteley et al., 2015; Robinson et al., 2020; Frankham, 2016, 2015). Theoretical studies have modelled the outcome of genetic rescue in specific situations to assess the risks and benefits to wild populations (Ash et al., 2023; Robinson et al., 2020). This allows for the exploration of the potential impact of different variables, such as inbreeding in the rescuing population (Kyriazis et al., 2021). There is also a growing body of experimental research testing how factors, such as the sex or degree of inbreeding in rescuers, and level of environmental stress, impact genetic rescue attempts (Zajitschek et al., 2009; Jørgensen et al., 2022; Heber et al., 2012; Hufbauer et al., 2015; Lewis et al., 2024). However, failures and negative effects have also been observed. For example, the Isle Royal wolf (*Canis lupus*) population collapsed following (naturally occurring) genetic rescue (Hedrick et al., 2014); in the Hihi (*Notiomystis cincta*) genetic rescue resulted in increased inbreeding 10 years later (Nichols et al., 2024); and in the Macquarie perch (*Macquaria australasica*) little or no mixing occurred between the rescuers and inbred population (Pavlova et al., 2024).

Despite the publication of guidelines as to when and where to attempt genetic rescue (Hedrick and Fredrickson, 2010; Frankham, 2015), there is still considerable reluctance by conservation stakeholders to attempt rescue in wild populations (Fitzpatrick et al., 2023). This is, to some degree, understandable due to potential risks such as outbreeding depression (Bell et al., 2019). This loss of fitness due to the crossing of two genetically divergent populations (Edmands, 2007) is associated with the breakdown of locally adapted gene complexes (Lenormand, 2002). An additional risk is genetic swamping, the rescuing population replacing unique genetic variation in the rescued population (Rhymer and Simberloff, 1996). Despite evidence to suggest such risks may be overstated, and that mixing divergent populations can provide considerable benefits (Kronenberger et al., 2017, 2018; Fitzpatrick et al., 2016), these risks highlight the importance of understanding what characterises the most effective rescuer(s) (Whiteley et al., 2015). Genetic structure of the rescuing population is an essential consideration (Kyriazis et al., 2021; Robinson et al., 2018; Ralls et al., 2020), as well as the number (van de Kerk et al., 2019; Kelly and Phillips, 2019) and the sex of rescuers (Zajitschek et al., 2009; Havird et al., 2016). These factors affect how much genetic diversity and load is introduced, how quickly it can introgress, and how long the rescue effect will last.

A central criticism of many genetic rescue studies is the fact that the longevity of rescue effects is not captured, due to the number of generations observed (Bell et al., 2019; Clarke et al., 2024). Laboratory studies on species with short generation times greatly facilitate our ability to monitor outcomes over multiple generations. Consequently, we can better test if and how quickly genetic rescue occurs, how long it lasts and whether there are any negative effects in the long term. In wild studies, where it is often extremely difficult and/or expensive to follow the rescue long-term, populations are often only monitored over a few consecutive generations (Hasselgren et al., 2018; Lotsander et al., 2021) or sporadically over generations (Miller et al., 2020).

Sex of rescuing individuals may be a key factor in the efficacy of genetic rescue as females are typically more limited in the number of offspring they can produce than males (Bateman, 1948). In many systems (i.e. promiscuous, polygynous, socially monogamous with extra-pair paternity), this means a male rescuer should speed the impact of genetic rescue. A male should sire more offspring carrying rescuing alleles and higher heterozygosity (Bateman, 1948) than a female, meaning that this additive variation is quicker to spread in the population. This effect has been shown in both guppies (*Poecilia reticulata*) (Zajitschek et al., 2009) and African lions (*Panthera leo*) (Miller et al., 2020; Trinkel et al., 2008). In addition, purging of genetic load is more effective in males due to differences in gamete investment between the sexes (Whitlock and Agrawal, 2009; Grieshop et al., 2021). In a rescue scenario, an individual with less genetic load should be favoured under sexual selection and have greater reproductive success.

Despite putative advantages of male rescue, female rescuers can be advantageous in other systems or scenarios. In the Florida panther (*Puma concolor couguar*), females were used for rescue (Pimm et al., 2006) as they were less likely to disperse or cause social conflict (Seal et al., 1994). Genetic load can also accumulate in mitochondrial DNA (mtDNA), which is commonly inherited through females (Gemmell et al., 2004). Thus, only female rescuers can introduce mtDNA variants to a population to reduce mtDNA genetic load (Gemmell and Allendorf, 2001). However, there is a risk of mitochondrial mismatch reducing offspring fitness (Havird et al., 2016). Female rescuers may also introduce maternal effects, the mother’s phenotype influencing that of the offspring (Wolf and Wade, 2009), which may affect rescue efficiency.

Another key consideration related to both rescuer sex and the genetic structure of rescuing populations is the background of sexual selection the rescuing population has experienced. Sexual selection can vary across populations (Kasumovic et al., 2008) affecting patterns of genetic variation (Parrett et al., 2022), facilitating adaptation (Parrett and Knell, 2018) and reducing inbreeding (Vega-Trejo et al., 2017). Stronger sexual selection has been shown to improve population fitness (Cally et al., 2019) and can also reduce genetic load in a population (Whitlock and Agrawal, 2009). Individuals from high sexual selection populations should also be more competitive in securing mates, thus gaining greater reproductive success, and increasing the speed at which genetic diversity introgresses during rescue. An increase in population fitness due to sexual selection has been observed in *Tribolium castaneum;* experimental populations experiencing elevated sexual selection were shown to be less likely to go extinct under stressful conditions than those that evolved under monogamy (Godwin et al., 2020; Lumley et al., 2015). Although beneficial, sexual selection may also promote assortative mating (van Doorn et al., 2009), and potentially reduce subsequent interbreeding between rescuers and rescued, thus hindering rescue attempts. To our knowledge, no studies have tested the effect of sexual selection background on the success of genetic rescue.

Here, we use the *Tribolium castaneum* model (Pointer et al., 2021) to experimentally address key omissions in the understanding of genetic rescue of inbred populations. *T. castaneum* has been utilised previously to study genetic rescue with one finding evidence of rescue (Hufbauer et al., 2015) and the other not observing a rescue effect (Lewis et al., 2024). First, we test if the sex of a rescuer has an impact on genetic rescue. We predict that a male rescuer will result in a greater fitness increase in inbred populations due to the reproductive difference between the sexes allowing faster introgression. Second, we test if rescuers evolved under different levels of sexual selection differentially impact the outcome of genetic rescue. We predict that a rescuer from a strong sexual selection background will be more effective, due to lowered genetic load. Importantly, we utilise the short generation time of *T. castaneum* to follow the effects of genetic rescue over 10 generations, allowing observation of both the speed and longevity of rescue effects. Additionally, we replicate our experimental populations under nutrient stress. We predict that stress will exaggerate the effects of inbreeding depression so that the magnitude of the rescue effect will be greater under stress than under benign conditions.

## Methods

### Ethics

No ethics approval was required for this study as experiments were conducted on an unregulated invertebrate species.

### Husbandry

*T. castaneum* were kept in a controlled environment at 30°C and 60% humidity with a 12:12 light-dark cycle. Populations were kept on standard fodder consisting of 90% organic white flour, 10% brewer’s yeast and a layer of oats for traction unless otherwise stated. During the husbandry cycle, 2mm and 850µm sieves were used to remove pupae and adults from fodder. The following cycle was started by a set number of adults (line dependent, see below) being placed into containers with fresh standard fodder. The oviposition phase: populations were given seven days to mate and lay eggs before adults were removed by sieving to prevent overlapping generations. The fodder containing eggs was returned to the container. The development phase: eggs were kept in the containers for 35 days to allow the eggs to develop into mature adults. Around day 21 of the development phase, pupae were collected to obtain known-sex virgin individuals which were then used to start the next generation. The pupae were kept as virgins in single-sex groups of 20 for 10 days to allow them to complete development. Once mature, the cycle began again with those beetles going into fresh fodder to form a population of males and females.

#### *Tribolium castaneum* lines

##### Krakow Super Strain (KSS)

was created by mixing fourteen laboratory strains to maximise genetic diversity in a single strain (Laskowski et al., 2015). This was used as the outbred treatment in the genetic rescue experiments.

##### Inbred Lines

Founded from KSS and inbred through three single-pair bottlenecks in the first, fifth and seventh generations. Between bottlenecks, the lines were maintained at a maximum population size of 100 randomly selected adults. Of the initial 30 lines, 24 survived the inbreeding treatment and 12 lines were maintained and used for experiments.

##### Sexual Selection Lines

polyandrous and monogamous lines were created from the Georgia 1 stock (Haliscak and Beeman, 1983; Lumley et al., 2015). Each polyandrous line (n=3) was maintained each generation in twelve groups each consisting of five males and one female. Following oviposition, the eggs from all groups in a line are mixed to form one population from which the next generation’s groups will be sourced. For each monogamous line (n=3) twenty separate mating pairs are bred. Following oviposition, the eggs from all pairs are mixed and the next pairs are sourced from this population to maintain that line. The number of groups and pairs in each regime results in a theoretical Ne = 40 in each treatment (Godwin et al., 2020). These regimes had been maintained for 150 generations when rescuers were taken. The polyandrous lines are hereafter referred to as sexual selection lines, and monogamous as no sexual selection.

#### Genetic rescue protocol

Replicate experimental inbred populations were created from the inbred lines to serve as populations to be rescued. Pupae were sexed and placed into plastic dishes with lids, containing 10ml standard fodder in single-sex groups. 10±2 days after eclosion, ten males and ten females from a given line were placed in a 125ml tub with 70ml of standard fodder creating populations each containing twenty adult beetles at a 1:1 sex ratio for the oviposition phase. On day 20±1 of the development phase, pupae were again taken from the populations using the method outlined above to create the next non-overlapping generation.

Populations were maintained using twenty reproducing adults per generation, not allowing population growth. This allowed us to maintain a roughly constant population density during offspring development across generations, avoiding the confounding influence of negative density-dependence on offspring production (Duval et al., 1939; King and Dawson, 1972; Janus, 1989).

Each experimental population was randomly assigned an ID number, to avoid bias when handling. After being established at the experimental size, the populations were maintained in experimental conditions for one generation to avoid transgenerational density effects affecting the genetic rescue results (Đukić et al., 2021). The rescue treatments were applied in the second generation under experimental conditions. In each population, a single beetle was replaced with a rescuer thus maintaining the 1:1 sex ratio and population size, avoiding any increase in productivity due to a demographic rescue. Rescuers taken from their source populations as pupae were age-matched as closely as possible to individuals in experimental populations. On day 37 of the development phase experimental populations were frozen at -6°C and mature offspring were counted as a measure of productivity (our metric for population fitness). If a population was removed from the experiment because of slow development (pupae were not available to establish the next generation), that population was analysed as part of all generations prior but excluded henceforth.

#### The sex of the rescuer in genetic rescue

Due to logistic issues with ventilation, four out of the 12 experimental inbred populations failed to produce offspring in generation 0. From each of the remaining eight inbred lines, three replicate populations were created and assigned to one of three treatments; No Rescue control (ten inbred line males, ten inbred line females); Male Rescue (nine inbred line males, one KSS male, ten inbred line females); and Female Rescue (ten inbred line males, nine inbred line females, one KSS female; Figure 1). Populations were maintained for ten, non-overlapping generations.

**Figure 1:**
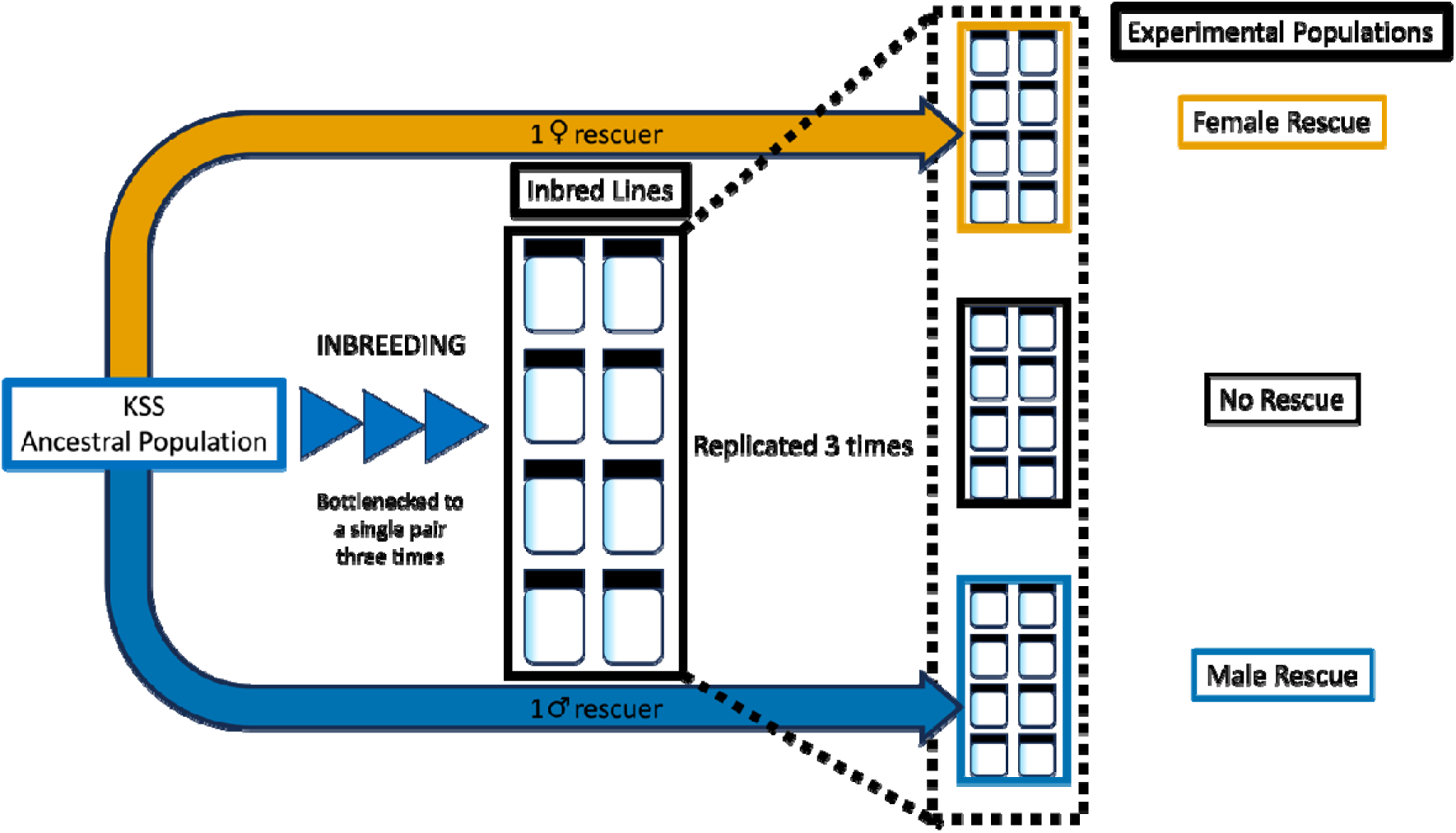
Experimental set-up of the creation and attempted genetic rescue of small, inbred *T. castaneum* populations (*N_e_* = 20) by a single male or female rescuer from the outbred ancestral population. Three experimental populations were created from each of 8 inbred lines resulting in 24 experimental populations, every line represented once in a treatment.

#### Sexual selection and genetic rescue

We investigated the impact of a rescuer’s sexual selection history on the effectiveness of genetic rescue. From 12 inbred lines, three replicate populations were created and assigned to one of three treatments; No Rescue Control (ten inbred line males, ten inbred line females); Sexual Selection Rescue (nine inbred line males, one polyandrous male and, ten inbred line females); No Sexual Selection Rescue (nine inbred line males, one monogamous male, ten inbred line females; Figure 2). A single polyandrous and single monandrous line were used as the source for rescuers. Populations were maintained for nine generations.

**Figure 2:**
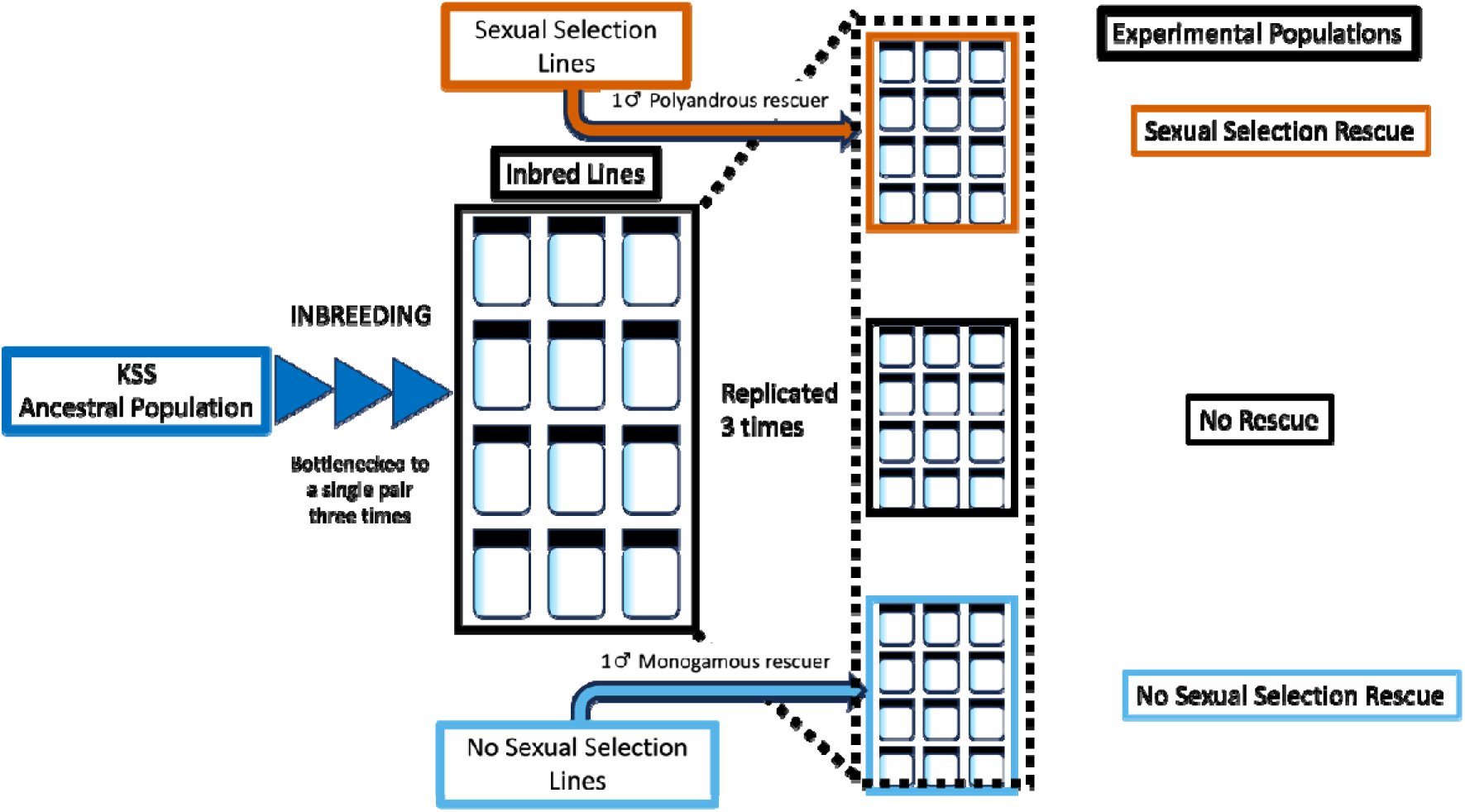
Experimental procedure for the creation and attempted genetic rescue of small, inbred *T. castaneum* populations (*N_e_* = 20) by a single male rescuer from either a sexual selection or no sexual selection line. Three experimental populations were created from each of 12 inbred lines resulting in 36 experimental populations, every line represented once in a treatment.

#### Stressful conditions

To test rescue under stress conditions, duplicate rescue populations were established from each rescued line at generation five in the ‘sex’ experiment, and generation six in the ‘sexual selection’ experiment. These were maintained as in the main experiments (until generation ten and nine respectively), but with a reduction in the yeast content of the fodder, which is the main source of protein for the experimental populations. This reduction generates nutrient stress in *T. castaneum* (Godwin et al., 2020). In the ‘sex’ experiment fodder contained 0% yeast and 1% yeast in the ‘sexual selection’ experiment (because of low survival with zero yeast).

#### Statistical analyses

Statistical analyses were carried out in R V4.4.1 (R Core Team, 2024) utilising R studio version 2024.04.2+764 (Posit team, 2024). Tidyverse (Wickham et al., 2019), stats (R Core Team, 2024), Rmisc (Hope, 2022)and googlesheets4 (Bryan, 2023) were used for data management and exploration. Plots were created using ggplot2 (Wickham, 2016). The distribution of data was checked using the shapiro.test function (R Core Team, 2024). Generalised Linear Mixed Models (GLMMs) were fitted to test for differences in productivity between the experimental treatments using glmmTMB (Brooks et al., 2017). Model fit was checked using DHARMa (Hartig, 2022). Model parameters were checked for collinearity using variance inflation factor (Vif) scores with the check_collinearity function from performance (Lüdecke et al., 2021). There were no issues with overdispersion or collinearity (VIF: <3 for all variables) in any models. R^2^ was determined using the r.squaredGLMM function in MuMIn (Bartoń, 2024). Post-hoc pairwise Tukey tests were carried out using multcomp (Hothorn et al., 2008).

Within each experiment, we fitted GLMMs with the same model structure, using a negative binomial distribution to model productivity counts, which provided better model fit than a Poisson distribution. Productivity was the response variable, with treatment, generation and generation^2^ as fixed effects. Inbred line of origin and experimental population ID were included as random effects, with ID nested within inbred line. Interaction terms (treatment x generation, treatment x generation^2^) were initially included but removed from the model if not significant. The generation^2^ factor was not significant in the models for populations under stressful conditions and was therefore removed. When a quadratic effect of generation was found in a model, we then tested to see if this was due to an increase and a decrease in productivity, rather than just one of these. We did this using two separate GLMMs (with the same factors as previously) run on the data split into generations 1-5 and 5-10. These GLMMs were fitted with treatment and generation as fixed effects, ID nested within inbred line as a random effect. GLMMs were also fitted on generations 2 and 3 individually (Table S3 + S4) in the ‘sex’ experiment, these single-generation models used a Poisson distribution, productivity as a response variable and treatment as a fixed effect. Random effects were the same as above. This was to test at which point the rescue treatments resulted in a significant difference from the control, to see if there were differences in the speed of male or female rescue.

## Results

### The sex of the rescuer in genetic rescue

Twenty-four populations were initiated, but in generation two one population in the control inbred populations failed to pupate in time for the next generation. That population was included in the analyses.

Male and female rescuer treatments both resulted in significantly higher productivity than the control (see Table 1, Figure 3). Generation^2^ also had a significant negative effect. Interactions between rescuer sex treatment x generation (and generation^2^) were not significant. In post-hoc tests there was no significant difference across all generations between the male and female rescue treatments (Estimate = 0.015, SE = 0.040, *z* = 0.374, *P* = 0.926, 95% CI = -0.079, -0.109).

**Figure 3:**
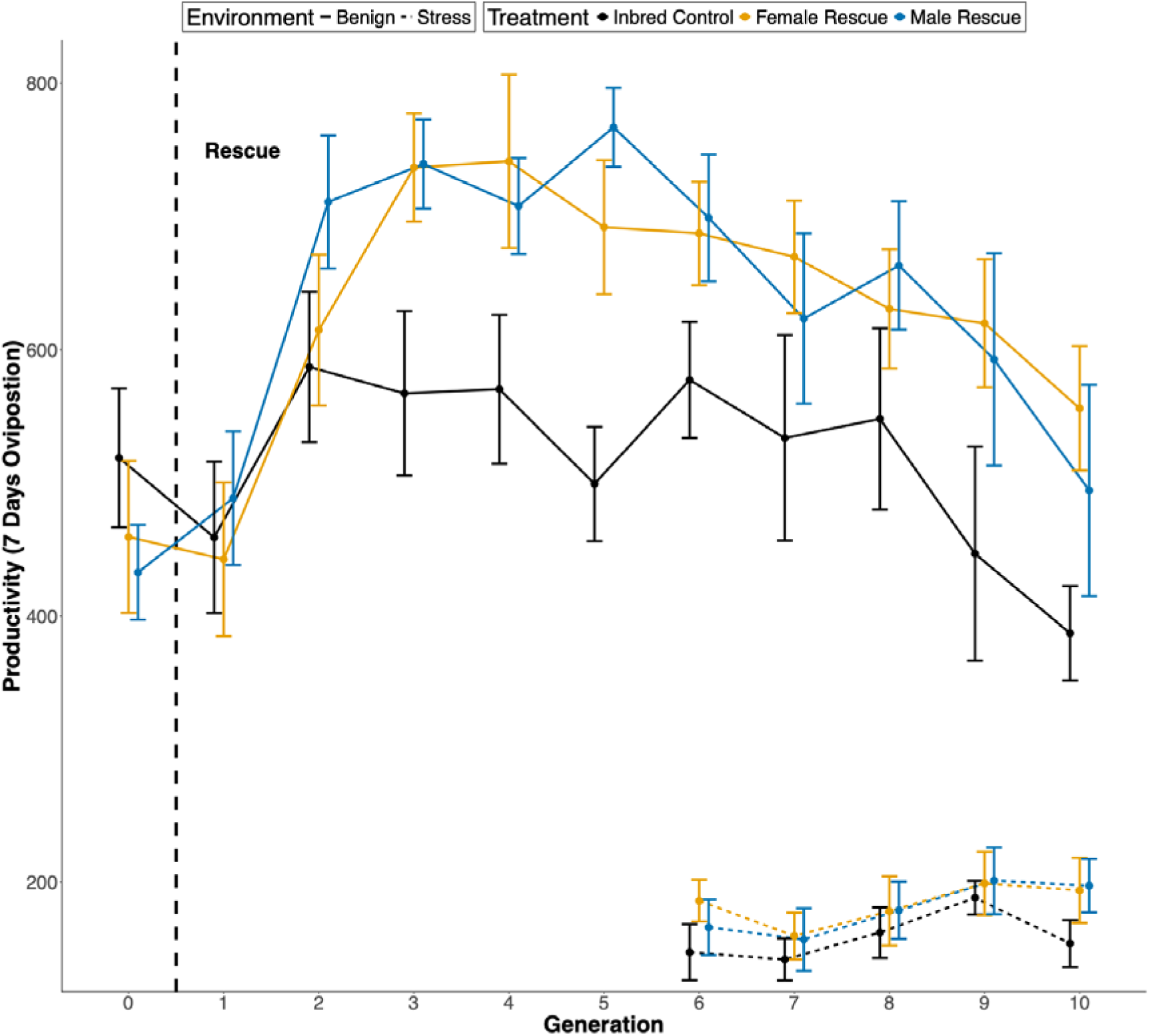
The effect of introducing a male or female rescuer on the mean productivity of small, inbred populations of *T. castaneum* (*N_e_* = 20, n = 24/23) over 10 generations after an introduction event. A single male or female rescuer was used to replace one individual of the same sex (dashed vertical line) within the populations of 10 females and 10 males. Populations were kept in either benign (solid line) or stressful (dashed line - starting only at generation 6) environmental conditions (fodder with or without yeast respectively). Under benign conditions, there was a significant increase in productivity for both male (Blue), and female (Orange) rescue treatments compared to the control treatment (Black). There was also a quadratic interaction with generation (See Table 1). Standard errors with 95% confidence intervals are shown.

**Table 1:** Factors impacting the productivity of small, inbred populations of *T. castaneum* (*N_e_* = 20, n = 24) receiving a single male or female genetic rescuer, or no rescue, tested using a GLMM.^1^ ^1^Productivity was measured over 10 generations following the rescue event. Predictors in bold are significant (*P <* 0.05). Marginal R^2^ = 0.247, Conditional R^2^ = 0.330. *One population was lost in Generation 2, so there are 23 populations from Generation 2 onwards.

| Predictor | Estimate | SE | z | P | 95% CI |
| --- | --- | --- | --- | --- | --- |
| Intercept | 5.944 | 0.075 | 79.570 | <2e-16 | 5.798<br>6.091 |
| Treatment (baseline = Control) |  |  |  |  |  |
| **<u>Female Rescue</u>** | **0.220** | **0.042** | **5.220** | **<0.001** | **0.138**<br>**0.303** |
| **<u>Male Rescue</u>** | **0.235** | **0.042** | **5.590** | **<0.001** | **0.153**<br>**0.318** |
| **<u>Generation</u>** | **0.160** | **0.027** | **6.030** | **<0.001** | **0.108**<br>**0.212** |
| **<u>Generation<sup>2</sup></u>** | **-0.015** | **0.002** | **-6.580** | **<0.001** | **-0.020**<br>**-0.011** |
| Random | 231 Observations |  | Variance |  |  |
| ID:Inbred line | 24 Populations* |  | 7.056e-10 |  |  |
| Inbred line | 8 Lines |  | 8.295e-03 |  |  |

Post-hoc tests showed that by generation two the productivity of the male rescue lines was significantly higher than the control lines (see Fig 1 & Table S3; Estimate = 0.189, SE = 0.092, *z* = 2.060, *P* = 0.040, 95% CI = 0.009, 0.370), but the productivity of the female rescue lines was not (see Fig 1 & Table S3; Estimate = 0.035, SE = 0.092, *z* = 0.370, *P* = 0.708, 95% CI = -0.146, 0.216). However, the productivity of male and female rescued lines in that generation (2) was not significantly different (see Fig 1; Estimate = 0.155, SE = 0.088, *z* = 1.760, *P* = 0.183, 95% CI = -0.051, 0.361). There was no significant difference between the male and female rescued lines in any other single generation (see Fig 1).

When modelled separately post-hoc, over generations 1-5 productivity increased significantly (see Table S1; Estimate = 0.070, SE = 0.016, *z* = 4.450, *P* < 0.001, 95% CI = 0.039, 0.101), then over generations 5-10 productivity decreased significantly (see Table S2; Estimate = -0.058, SE = 0.014, *z* = -4.26, *P* < 0.001, 95% CI = -0.085, -0.031).

Under stress conditions (0% yeast in fodder) productivity greatly decreased (Figure 3), and there were no significant differences between the treatments. There was a significant linear effect of generation on productivity (see Table 2).

**Table 2:** Factors impacting the productivity of small, inbred *T. castaneum* populations (*N_e_* = 20, n = 23) under nutrient stress that had either a male or female rescuer from an outbred population introduced five generations prior, tested using a GLMM.^2^ ^2^Predictors in bold are significant (*P* < 0.05).

| Predictor | Estimate | SE | *z* | *P* | 95%<br>CI |
| --- | --- | --- | --- | --- | --- |
| Intercept | 4.716 | 0.136 | 34.570 | <2e-16 | 4.449<br>4.984 |
| Treatment (baseline = Control) |  |  |  |  |  |
| Female Rescue | 0.221 | 0.117 | 1.890 | 0.059 | -0.008<br>0.451 |
| Male Rescue | 0.113 | 0.118 | 1.040 | 0.300 | -0.109<br>0.354 |
| **<u>Generation</u>** | **0.038** | **0.012** | **3.190** | **0.001** | **0.015**<br>**0.062** |
| Random | 78 Observations |  | Variance |  |  |
| ID:Inbred line | 23 Populations |  | 0.040 |  |  |
| Inbred line | 8 Lines |  | 0.021 |  |  |

<sup>2</sup> Predictors in bold are significant ( $P < 0.05$ ).

### Sexual Selection and Genetic Rescue

Thirty-six populations were initiated, but in both generations two and five one population in the control inbred populations failed to pupate in time for the next generation. These populations were included in the analyses.

Introducing a rescuer from a sexual selection population had a significant negative effect on productivity that interacted with generation^2^; i.e., there was a significant increase in productivity between generation 2-5, after which productivity declined. There was no evidence of an interaction between ‘no sexual selection’ rescue and generation^2^ (Figure 4, Table 3). There was no significant effect when the interaction was removed.

**Figure 4:**
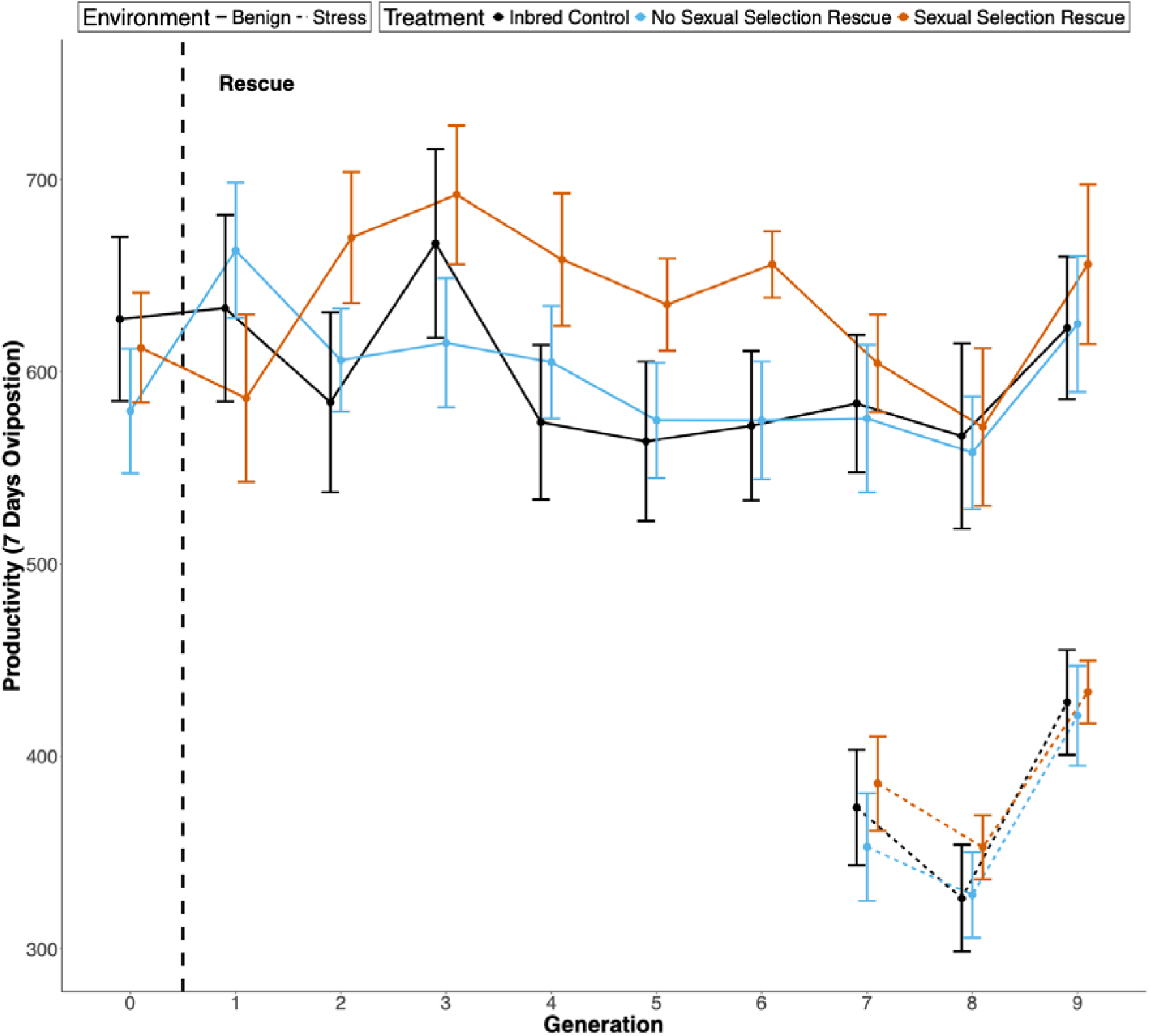
The effect of introducing a single male genetic rescuer from a sexual selection background or no sexual selection background on the productivity of small, inbred *T. castaneum* populations (*N_e_* = 20, n = 36/34) over nine generations. Populations were in either a benign (solid line) or stressful (dashed line) environment. The rescue was a single event (dashed vertical line) where the rescuer replaced a male in the inbred population. Compared to the control (black) there was a significant increase in productivity in the sexual selection rescue treatment (orange), which had a quadratic interaction with generation, but no significant effect of the no sexual selection treatment (blue) (See Table 2). Standard errors with 95% confidence intervals are shown.

**Table 3:**
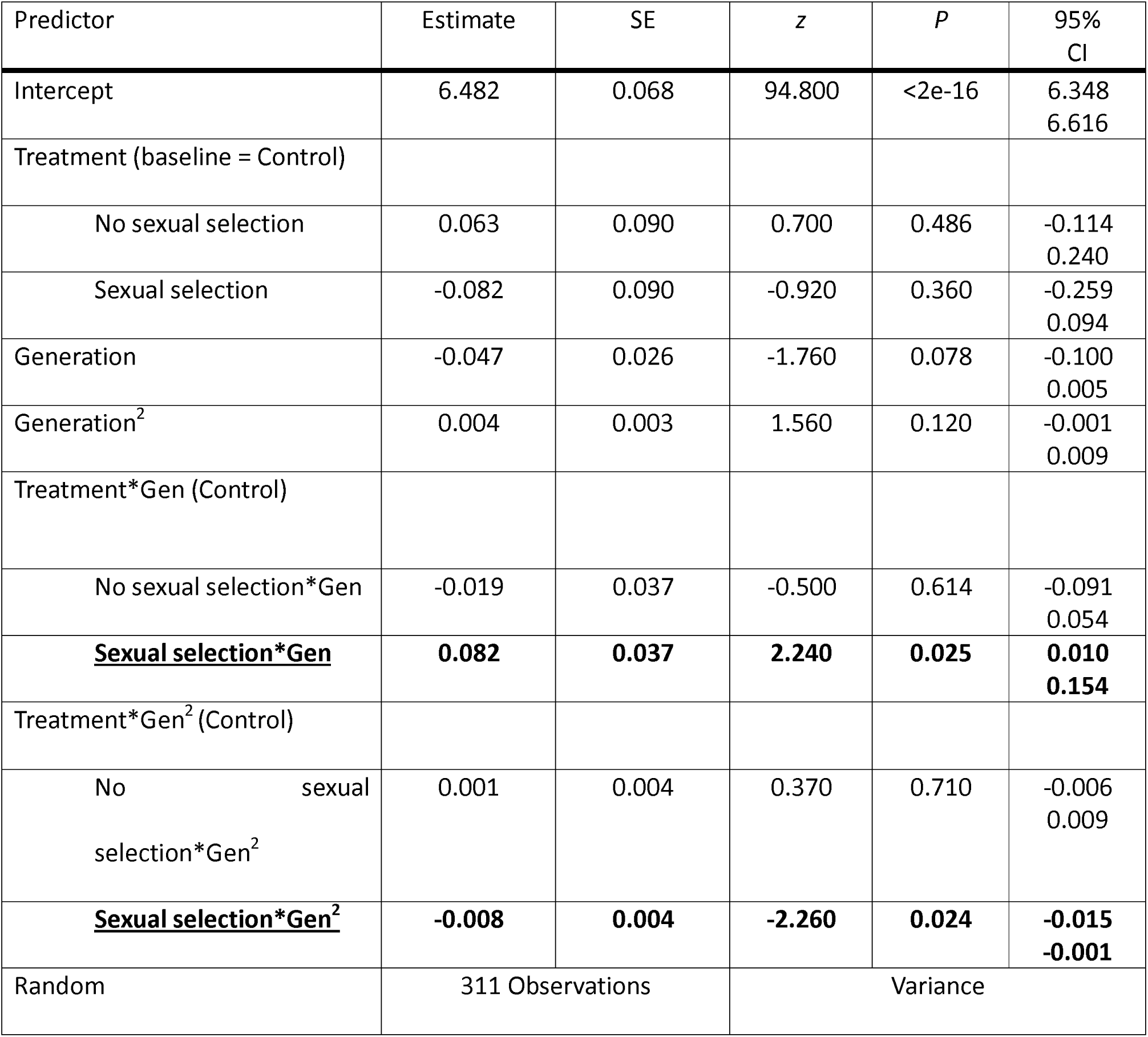
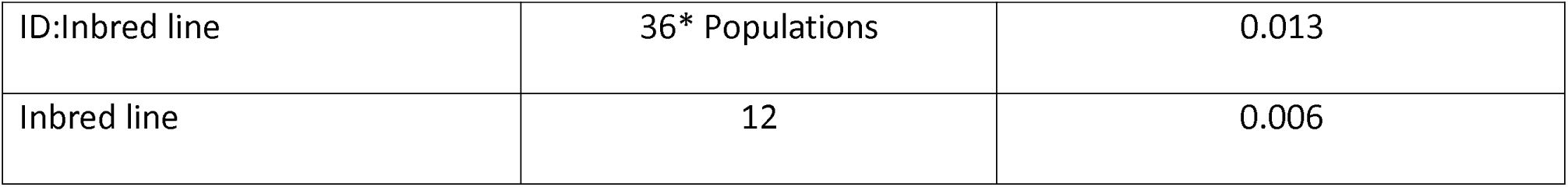
Factors impacting the productivity of small, inbred populations (*N_e_* = 20, n = 36) of *T. castaneum* that received a single rescuer from either a sexual selection or no sexual selection background line population, tested using a GLMM.^3^ ^3^Predictors in bold are significant (*P* < 0.05). Marginal R^2^ = 0.077, Conditional R^2^ = 0.512. *One population was lost in Generation 2 and one in Generation 5.

| Predictor | Estimate | SE | z | P | 95% CI |
| --- | --- | --- | --- | --- | --- |
| Intercept | 6.482 | 0.068 | 94.800 | <2e-16 | 6.348<br>6.616 |
| Treatment (baseline = Control) |  |  |  |  |  |
| No sexual selection | 0.063 | 0.090 | 0.700 | 0.486 | -0.114<br>0.240 |
| Sexual selection | -0.082 | 0.090 | -0.920 | 0.360 | -0.259<br>0.094 |
| Generation | -0.047 | 0.026 | -1.760 | 0.078 | -0.100<br>0.005 |
| Generation<sup>2</sup> | 0.004 | 0.003 | 1.560 | 0.120 | -0.001<br>0.009 |
| Treatment*Gen (Control) |  |  |  |  |  |
| No sexual selection*Gen | -0.019 | 0.037 | -0.500 | 0.614 | -0.091<br>0.054 |
| **<u>Sexual selection\*Gen</u>** | **0.082** | **0.037** | **2.240** | **0.025** | **0.010**<br>**0.154** |
| Treatment*Gen<sup>2</sup> (Control) |  |  |  |  |  |
| No sexual selection*Gen<sup>2</sup> | 0.001 | 0.004 | 0.370 | 0.710 | -0.006<br>0.009 |
| **<u>Sexual selection\*Gen<sup>2</sup></u>** | **-0.008** | **0.004** | **-2.260** | **0.024** | **-0.015**<br>**-0.001** |
| Random | 311 Observations |  | Variance |  |  |

<sup>3</sup> Predictors in bold are significant ( $P < 0.05$ ). Marginal $R^2 = 0.077$ , Conditional $R^2 = 0.512$ . \*One population was lost in Generation 2 and one in Generation 5.

|  |  |  |
| --- | --- | --- |
| ID:Inbred line | 36* Populations | 0.013 |
| Inbred line | 12 | 0.006 |

Under stress conditions, there were no significant differences between the treatments’ productivity, but productivity did increase over generations (Figure 2, Table 4).

**Table 4:** Factors impacting the productivity of small, inbred populations (*N_e_* = 20, n = 34) of *T. castaneum* under nutrient stress that had been rescued by either a high or low sexual selection male rescuer seven generations prior, tested using a GLMM.^4^ ^4^Predictors in bold are significant (*P* < 0.05).

| Predictor | Estimate | SE | *z* | *P* | 95%<br>CI |
| --- | --- | --- | --- | --- | --- |
| Intercept | 5.281 | 0.180 | 29.385 | <2e-16 | 4.929<br>5.633 |
| Treatment (baseline = Control) |  |  |  |  |  |
| No sexual selection Rescue | -0.023 | 0.066 | -0.352 | 0.725 | -0.152<br>0.106 |
| Sexual selection Rescue | 0.048 | 0.065 | 0.741 | 0.459 | -0.080<br>0.177 |
| **<u>Generation</u>** | **0.079** | **0.021** | **3.731** | **<0.001** | **0.038**<br>**0.120** |
| Random | 102 Observations |  | Variance |  |  |
| ID:Inbred line | 34 Populations |  | 0.013 |  |  |
| Inbred line | 12 |  | 0.011 |  |  |

## Discussion

We tested how the sex and sexual selection evolutionary history of a rescuing individual affects the duration of genetic rescue using small, inbred populations of *T. castaneum*. Our results show that a male or a female rescuer was equally effective; both improved productivity compared to the control, though there was some evidence that a male rescuer led to faster rescue. In the second experiment, the introduction of a male from an elevated sexual selection background resulted in a significant increase in productivity, whilst a male from a monogamous background did not. Importantly, in both experiments we observed temporal effects; in the successful rescue treatments productivity increases were observed in the initial generations after the introduction of rescuers, before declining in later generations. When these experiments were replicated under severe nutrient stress conditions we saw no significant effect of rescue on productivity.

Male rescuers have been suggested to enable faster/greater genetic rescue than females due to their higher reproductive potential, as generating more offspring will spread introduced genetic diversity faster (Zajitschek et al., 2009). Our results suggest that females are as effective at rescuing the inbred populations as males. We did find some evidence that males may enable faster rescue of productivity; with male rescue lines showing a significantly earlier increase in productivity compared to control lines (by generation 2) than female (by generation 3) rescue lines (See Figure 1 & Table S3). This did not translate into a significant difference between the productivity of male and female rescue lines in generation 2. This result contrasts with previous studies: in wild lions, males were more effective rescuers despite potential issues of social disruption and infanticide (Trinkel et al., 2008), and in guppies, faster population growth was observed following male rescue (Zajitschek et al., 2009). One aspect which may explain these differences is the extreme disparity between the mating systems of target species, coupled with our experimental approach. We used smaller populations (*N_e_* = 20) than in other studies of genetic rescue in *T. castaneum* (Durkee et al., 2023; Hufbauer et al., 2015; Lewis et al., 2024), which may have limited the advantage that male rescuers had over female rescuers. As female *T. castaneum* can mate with 4-6 males in an hour (Pai and Yan, 2003), the 10 females available to a male in our populations over seven days is far less than his mating potential, and thus the impact of genetic rescue. More experimentation is needed, factoring in population size and testing species with different variation in reproductive success between sexes.

*T. castaneum* is a promiscuous and highly fecund species (Pointer et al., 2021) and our results are applicable to species with similar life history strategies and mating systems. Females in this system may act as equivalent rescuers to males as there is evidence of inbreeding avoidance in the female reproductive behaviour (Attia and Tregenza, 2004; Michalczyk et al., 2011) meaning negative impacts of inbreeding (Hedrick and Garcia-Dorado, 2016; Vega-Trejo et al., 2022) may be minimised. However, *T. castaneum* females do not exhibit care for offspring (Kölliker, 2012), eliminating a potential advantage provided by a female rescuer (Mattey et al., 2018; Pooley et al., 2014).

We predicted that rescuers drawn from populations with elevated sexual selection would be more fit (with less genetic load) and more competitive, resulting in a more effective genetic rescue. Our results support this, rescuers with a high sexual selection background improved productivity in the inbred populations where rescuers from a no sexual selection background did not. The lines from which our sexually selected rescuers were sourced have previously been shown to resist extinction in the face of inbreeding, relative to lines with no history of sexual selection (Godwin et al., 2020; Lumley et al., 2015). This suggests that these lines have a higher fitness due to sexual selection. Using males from these lines as rescuers may have increased productivity for several reasons, including increased mating competitiveness and increased fitness in offspring with lower genetic load. Furthermore, lower introduced genetic load should result in less re-emergent inbreeding depression in later generations in these small populations. Further work is needed to unravel these possibilities

The effects of inbreeding depression on endangered populations are often exacerbated by exposure to environmental stress (Armbruster and Reed, 2005; Richardson et al., 2004). However, when testing rescue treatments under stressful (nutrient) conditions we found no significant differences between treatments in either sex or sexual selection experiments. This was unexpected as stress should magnify inbreeding depression and disproportionally affect the productivity of populations that had not been rescued. This lack of effect may be due to the harshness of the nutrient stress treatment we used, as this has been shown to greatly reduce female fecundity and slow offspring development (Demont et al., 2014). Nutrient stress could also increase cannibalism, which occurs in *T. castaneum* when food is scarce (King and Dawson, 1972). This may have had more impact on rescued populations due to increased competition for resources when initial productivity (eggs laid) is higher. However, stressful conditions do not always exaggerate inbreeding depression (Armbruster and Reed, 2005; Sandner et al., 2022). Our finding that stress repressed genetic rescue points to the importance of improving environmental conditions for species before attempting to recover population numbers (Bell et al., 2019; Root, 1998; Ferreras et al., 2001).

A regular criticism of genetic rescue studies is that they fail to monitor populations over sufficient timescales (Clarke et al., 2024). Our study continued monitoring rescue outcomes over multiple (9-10) generations. We see genetic rescue effects begin in the second generation after rescue. Rescue effects are not seen in the generation immediately following rescue, likely because, even in a promiscuous population, it will take more than one generation for the variation from a single rescuer to introgress widely into the population and influence overall productivity. In both experiments, the treatments that result in rescue have peak productivity around generation 5-6. This suggests the beneficial introgression of the rescuer’s genetic diversity into the population takes several generations, as seen in previous studies (Hufbauer et al., 2015).

Importantly, we saw productivity benefits of rescue began to decline by the sixth generation in both experiments. Many genetic rescue studies are relatively short-term projects relative to the generation time of the species involved (Frankham, 2015). Owing to the short generation times of *T. castaneum* (Pointer et al. 2021), we are the first to show that rescue effects may be short-lasting. This has important connotations for studies in wild systems, reinforcing suggestions that monitoring must continue in the long-term, but also that single rescue introductions are potentially not sufficient to rescue populations. We suggest our findings are associated with the resumption of inbreeding effects in later generations due to small population size (N = 20). This does not reduce the relevancy to conservation contexts, as similar effects have been seen in wild systems (Hedrick et al., 2014, 2019; Robinson et al., 2019). The genetic rescue of the Florida Panther resulted in benefits for five generations after rescue (Onorato et al., 2024), our results suggest that in the coming generations these benefits may start to decline.

In conclusion, we find that both male and female rescuers can be effective genetic rescuers. This is likely linked to the dynamics of promiscuous mating systems such as that seen in *T. castaneum* but serves to highlight the importance of such species-dependent traits when planning conservation interventions. Importantly, and for the first time, we show sexual selection background affects the efficacy of genetic rescue. Given these results, we suggest that, where feasible, using a rescuer from a high sexual selection background when attempting genetic rescue could be beneficial in conservation programs. Overall, our results add important evidence to our understanding of the effectiveness of genetic rescue and support the argument that it should be considered an important tool to conserve endangered populations.

## Acknowledgements

We thank the members of the UEA *Tribolium* Lab for assistance with line maintenance and data collection.

