## Supplemental Tables for "Sexual selection matters in genetic rescue, but productivity benefits fade over time; a multi-generation experiment to inform conservation"

Supporting

Table S1: Factors impacting the productivity of small, inbred populations (*N_e_* = 20, n = 24) of *T. castaneum* rescued by either a male or female rescuer in the first five generations following rescue. Tested using a GLMM. Predictors in bold are significant (*P* < 0.05).

| Predictor | Estimate | SE | *z* | *P* | 95%  CI |
| --- | --- | --- | --- | --- | --- |
| Intercept | 6.054 | 0.073 | 82.800 | <2e-16 | 5.911  6.197 |
| Treatment (Control) |  |  |  |  |  |
| **Female Rescue** | **0.191** | **0.054** | **3.550** | **<0.001** | **0.086**  **0.296** |
| **Male Rescue** | **0.255** | **0.054** | **4.760** | **<0.001** | **0.150**  **0.360** |
| **Generation** | **0.070** | **0.016** | **4.450** | **<0.001** | **0.039**  **0.101** |
| Random | 116 Observations | | Variance | | |
| ID:Inbred line | 24 Populations | | <0.001 | | |
| Inbred line | 8 Lines | | <0.001 | | |

Table S2: Factors impacting the productivity of small, inbred populations (*N_e_* = 20, n = 24) of *T. castaneum* rescued by either a male or female rescuer in generations five to ten following rescue. Tested using a GLMM. Predictors in bold are significant (*P* < 0.05).

| Predictor | Estimate | SE | *z* | *P* | 95%  CI |
| --- | --- | --- | --- | --- | --- |
| Intercept | 6.638 | 0.113 | 58.890 | <2e-16 | 6.417  6.858 |
| Treatment (Control) |  |  |  |  |  |
| **Female Rescue** | **0.259** | **0.057** | **4.530** | **<0.001** | **0.147**  **0.371** |
| **Male Rescue** | **0.250** | **0.057** | **4.370** | **<0.001** | **0.138**  **0.363** |
| **Generation** | **-0.058** | **0.014** | **-4.260** | **<0.001** | **-0.085**  **-0.031** |
| Random | 116 Observations | | Variance | | |
| ID:Inbred line | 23 Populations | | <0.001 | | |
| Inbred line | 8 Lines | | <0.001 | | |

Table S3: Factors impacting the productivity of small, inbred populations (*N_e_* = 20, n = 24) of *T. castaneum* that had been rescued by either a male or female rescuer in the second generation following rescue. Tested using a GLMM. Predictors in bold are significant (*P* < 0.05).

| Predictor | Estimate | SE | *z* | *P* | 95%  CI |
| --- | --- | --- | --- | --- | --- |
| Intercept | 6.360 | 0.083 | 76.550 | <2e-16 | 6.198  6.523 |
| Treatment (Control) |  |  |  |  |  |
| Female Rescue | 0.035 | 0.092 | 0.370 | 0.708 | -0.146  0.216 |
| **Male Rescue** | **0.189** | **0.092** | **2.060** | **0.040** | **0.009**  **0.370** |
| Random | 23 Observations | | Variance | | |
| ID:Inbred line | 23 Populations | | 0.171 | | |
| Inbred line | 8 Lines | | 0.135 | | |

Table S4: Factors impacting the productivity of small, inbred populations (*N_e_* = 20, n = 24) of *T. castaneum* rescued by either a male or female rescuer in the third generation following rescue. Tested using a GLMM. Predictors in bold are significant (*P* < 0.05).

| Predictor | Estimate | SE | *z* | *P* | 95%  CI |
| --- | --- | --- | --- | --- | --- |
| Intercept | 6.290 | 0.073 | 86.42 | <2e-16 | 6.147  6.432 |
| Treatment (Control) |  |  |  |  |  |
| **Female Rescue** | **0.302** | **0.077** | **3.900** | **<0.001** | **0.150**  **0.454** |
| **Male Rescue** | **0.309** | **0.077** | **3.990** | **<0.001** | **0.157**  **0.460** |
| Random | 23 Observations | | Variance | | |
| ID:Inbred line | 23 Populations | | 0.171 | | |
| Inbred line | 8 Lines | | 0.135 | | |
